# CRIMSON: An Open-Source Software Framework for Cardiovascular Integrated Modelling and Simulation

**DOI:** 10.1101/2020.10.14.339960

**Authors:** C.J. Arthurs, R. Khlebnikov, A. Melville, M. Marčan, A. Gomez, D. Dillon-Murphy, F. Cuomo, M.S. Vieira, J. Schollenberger, S.R. Lynch, C. Tossas-Betancourt, K. Iyer, S. Hopper, E. Livingston, P. Youssefi, A. Noorani, S. Ben Ahmed, F.J.H. Nauta, T.M.J. van Bakel, Y. Ahmed, P.A.J. van Bakel, J. Mynard, P. Di Achille, H. Gharahi, K. D. Lau, V. Filonova, M. Aguirre, N. Nama, N. Xiao, S. Baek, K. Garikipati, O. Sahni, D. Nordsletten, C.A. Figueroa

## Abstract

In this work, we describe the CRIMSON (CardiovasculaR Integrated Modelling and SimulatiON) software environment. CRIMSON provides a powerful, customizable and user-friendly system for performing three-dimensional and reduced-order computational haemodynamics studies via a pipeline which involves: 1) segmenting vascular structures from medical images; 2) constructing analytic arterial and venous geometric models; 3) performing finite element mesh generation; 4) designing, and 5) applying boundary conditions; 6) running incompressible Navier-Stokes simulations of blood flow with fluid-structure interaction capabilities; and 7) post-processing and visualizing the results, including velocity, pressure and wall shear stress fields. A key aim of CRIMSON is to create a software environment that makes powerful computational haemodynamics tools accessible to a wide audience, including clinicians and students, both within our research laboratories and throughout the community. The overall philosophy is to leverage best-in-class open source standards for medical image processing, parallel flow computation, geometric solid modelling, data assimilation, and mesh generation. It is actively used by researchers in Europe, North and South America, Asia, and Australia. It has been applied to numerous clinical problems; we illustrate applications of CRIMSON to real-world problems using examples ranging from pre-operative surgical planning to medical device design optimization. CRIMSON binaries for Microsoft Windows 10, documentation and example input files are freely available for download from www.crimson.software, and the source code with compilation instructions is available on GitHub https://github.com/carthurs/CRIMSONFlowsolver (CRIMSON Flowsolver) under the GPL v3.0 license, and https://github.com/carthurs/CRIMSONGUI (CRIMSON GUI), under the AGPL v3.0 license. Support is available on the CRIMSON Google Groups forum, located at https://groups.google.com/forum/#!forum/crimson-users.

## Introduction

One of the revolutionary successes of twentieth century in applied mathematics is the development of the finite element method (FEM) into a reliable engineering tool. It is now routinely deployed by practitioners in fields where the problems of interest are described by systems of partial differential equations (PDEs). FEM, in combination with high-performance computing (HPC), has enabled solution of these systems of PDEs rapidly and accurately.

In the present work, FEM is applied to the incompressible Navier-Stokes equations in patient-specific vascular geometries. This enables analysis of clinically-relevant blood flow phenomena including the impact of the geometry, vascular wall properties, or prospective surgical decisions on blood pressure, wall shear stress, and mass (e.g. protein or drug) transport. These capabilities support the key applications of surgical planning, patient diagnosis, medical device design and optimisation, and basic cardiovascular disease research. Such applications are collectively called *computational haemodynamics* (CH).

A serious factor limiting widespread adoption of CH has been the arcane and unwieldy nature of existing academic software packages and workflows, coupled with the fact that no commercial package truly supports cutting-edge CH. Due in part to the patchwork nature of academic software development, involving multiple researchers with differing goals over many years, simulation workflows often require using multiple pieces of software and ad-hoc scripts. One pathway to addressing this issue is specialised, problem-specific commercial enterprise. For example, HeartFlow Ltd. provides CH-as-a-service for diagnosis of coronary artery disease, taking medical imaging data and producing computational clinical reports on the hemodynamic significance of coronary lesions. In their workflow, medical image data are uploaded to the HeartFlow servers, and a report is returned to the clinician (Taylor, et al., 2013). However, if CH is to be performed in a research setting using open-source tools, custom workflows must be created. Given the complexity of the software components and the interdisciplinary nature of the user base (e.g., clinicians, students, and academics), it is important that the software provides powerful capabilities, whilst presenting an intuitive, expandable, and user-friendly graphical user interface (GUI). In this article, we describe how we address these issues in our CH software environment CRIMSON (<u>C</u>ardiovascula<u>R</u> Integrated <u>M</u>odelling and <u>S</u>imulati<u>ON</u>).

CRIMSON consists of two components. The first is the CRIMSON GUI, which is a Windows application for setting up simulations via segmenting medical image data, creating finite element meshes, assigning boundary conditions, and examining results. The second component is the CRIMSON Flowsolver, which is a Message Passing Interface (MPI) parallel application, scalable to tens of thousands of cores (Zhou, et al., 2010), which can be run on Windows or Linux; simulations with the Flowsolver can either be run on the local machine from within the GUI, or transferred to a high-performance computer for simulation there.

The conceptual ancestor of CRIMSON is SimVascular (Updegrove, et al., 2016), which has been widely used in numerous CH studies (Sengupta, et al., 2012) (Marsden, et al., 2009) (Marsden, et al., 2013) (Les, et al., 2010) (Coogan, et al., 2013). During the early design phase of CRIMSON, we determined that there was a pressing need for developing a CH software package that leveraged the strengths of SimVascular’s Flowsolver with a modern UI with broad community support and best-in-class open source components for key operations of the simulation workflow (CAD, boundary condition specification, meshing, parameter estimation, etc.). For example, assigning a boundary condition directly to a surface of the 3D-rendered model was not supported by SimVascular’s GUI. CRIMSON GUI was therefore created to avoid these limitations, introduce many novel features and improvements, and expand functionality, workflow robustness, and accessibility. (). A key decision was to utilise for the first time in CH (Khlebnikov and Figueroa, 2016) the Medical Imaging Interaction Toolkit (MITK) (Wolf, et al., 2005). CRIMSON allowed the creation of novel and unique features including GUI support for a wide range of fine-tuneable boundary condition (BC) types, including direct imposition of phase-contrast magnetic resonance imaging (PC-MRI) flow data (Gomez et al. 2018), rapid design and prototyping of arbitrary lumped parameter network (LPN) reduced order models, and spatially-varying vessel wall material properties, all of which are supported in a modern, user-friendly GUI. Furthermore, CRIMSON supports automatic model parameter estimation via filtering of time-resolved data (Arthurs, et al., 2020). The key driving philosophy is that for these powerful features to reach their maximum potential, they must be accessible to both clinicians and technical researchers. To achieve this, CRIMSON aims to present clearly-documented, logical workflows in the GUI, and to protect users from common errors. The latter goal is achieved, for example, by supporting clear graphical BC specification in the GUI, by including undo/redo stacks for operations, and by limiting the interaction that users have with text-based configuration files.

Beyond making the basic workflow as user-friendly as possible, CRIMSON has several unique features which enable novel simulation capabilities. For example, CRIMSON allows users to implement – at run-time – arbitrary rules for adjusting boundary condition parameters during simulations, based upon the simulation state. This allows modelling of cardiovascular control mechanisms, and transitional hemodynamic states – for example, between rest and exercise (Arthurs, Lau et al., 2016). Further, for model parameterisation, testing, and exploring long-term behaviour, CRIMSON supports a one-click-to-enable pure-zero-dimensional mode, so that the haemodynamic influence of the boundary conditions can be explored without the computational cost of a full 3D domain. This is invaluable for novices, and for experts working to understand complex models.

## Design and Implementation

The CRIMSON workflow, from medical imaging data to simulation results, is shown in Figure 1. The three major stages of the pipeline (i.e., Preprocessing, Flowsolver, and Postprocessing) are depicted, including some optional sub-steps in each column which may or may not be part of the process for a given CH application.

**Figure 1.**
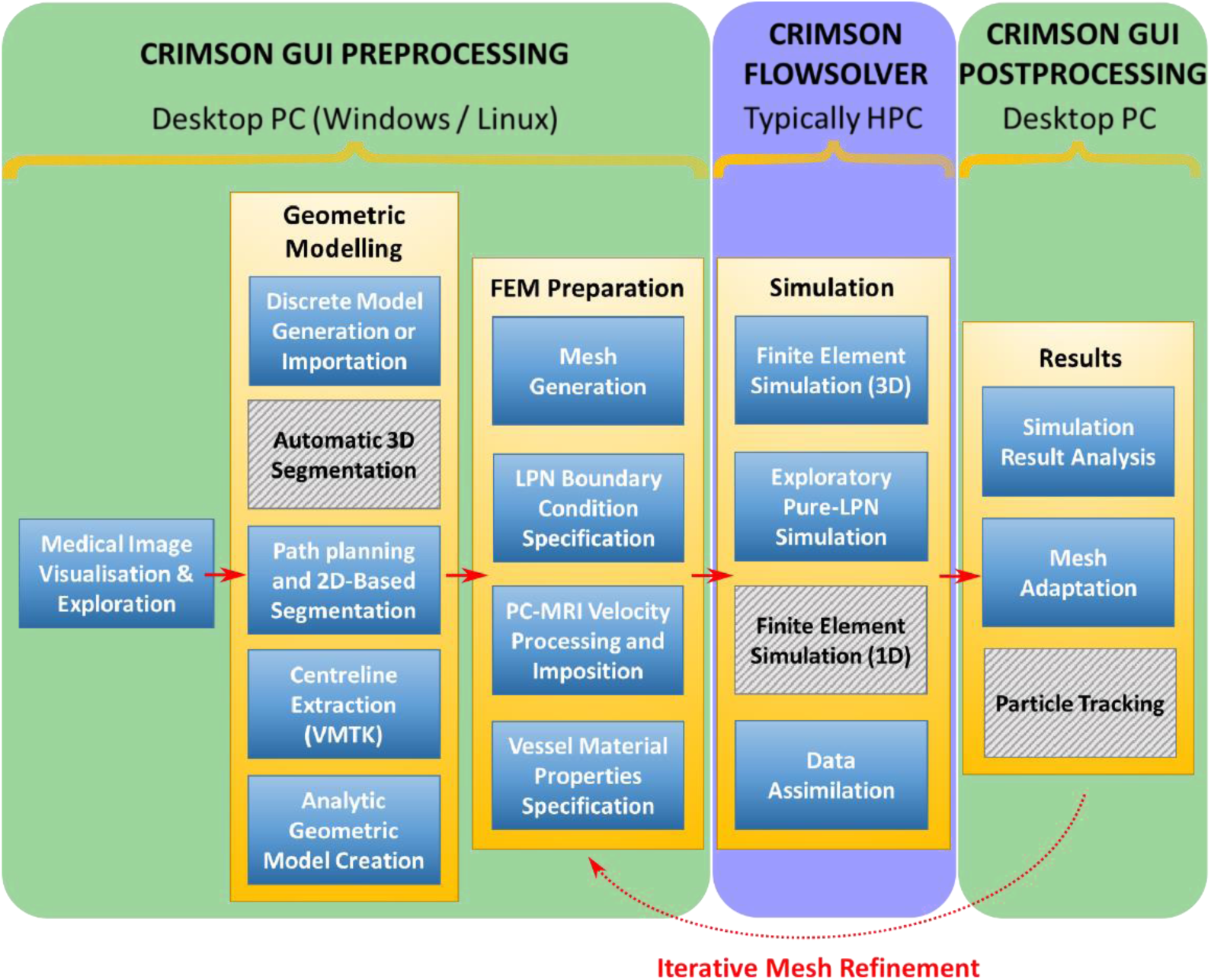
The CRIMSON workflow consists of Preprocessing, Flowsolver, and Postprocessing stages represented in three different columns. Typical hardware architecture utilized at each stage is noted. Some of the steps within each stage are optional, depending upon the complexity of the simulation. Grey hashed boxes indicate areas where the GUI support is under development. A medical image volume is required as initial input, if not importing a discrete geometric model from elsewhere. The final output is, at minimum, time-resolved pressure and velocity fields throughout the domain.

The CRIMSON GUI is built upon MITK, which provides tools to load, visualize and manipulate medical image data. A screenshot of CRIMSON during the Geometric Modelling portion of the workflow is shown in Figure 2. Here, the user loads the medical image volume, and draws one-dimensional paths approximately representing the centreline of each vessel of interest. CRIMSON then re-slices the image data using a plane perpendicular to the centreline at each point along its length (Figure 2, box 3). Upon this re-slice plane, the user generates two-dimensional contours to segment the vessel wall (Wang, et al., 1999) (Wilson, et al., 2001) using MITK’s manual and semi-automatic segmentation functions, including tools for delineating the vessel cross-section using circles, ellipses or splines. An analytic NURBS surface is then created of each vessel via lofting or sweeping operations. The vessels of interest are united using a blending operation to create a single geometric CAD model (Open Cascade, 2016), which is then volume-meshed using linear tetrahedral elements. Mesh generation is provided by TetGen (Si, 2015). CAD models can be exported in step, iges, and brep file formats. CRIMSON also allows for direct importing of discrete surface triangulations of vascular geometries (e.g. stl format) created in external packages.

**Figure 2:**
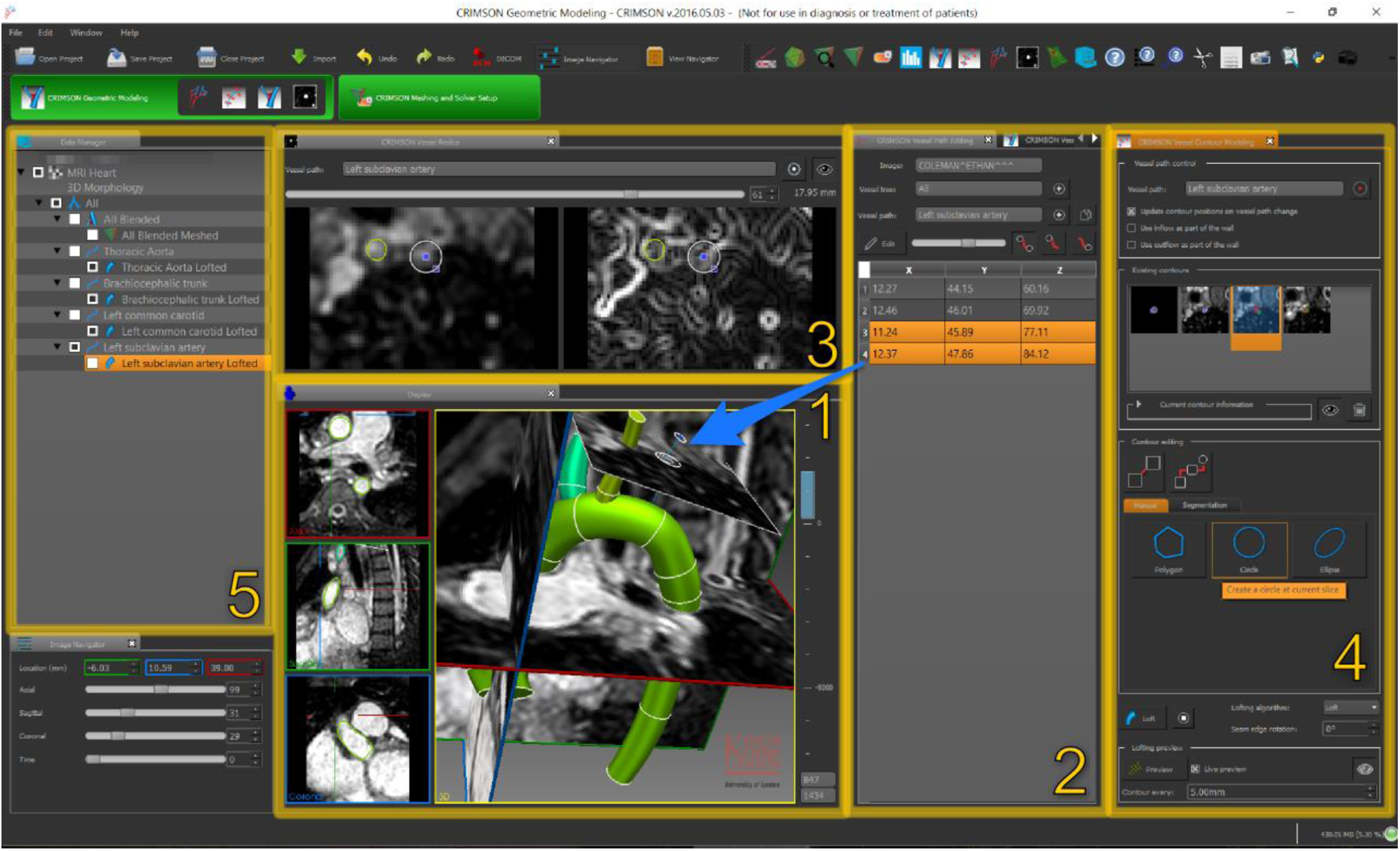
Geometric modelling aspect of the CRIMSON GUI. (1) The image data and segmentation is viewed in three adjustable orthogonal planes and a 3D projection. (2) Centerlines are created for each vessel of interest, editable via coordinates in Panel 2, or 3D interaction in Panel 1 (blue arrow shows relationship). (3) A “re-slice” of the image, consisting of a plane perpendicular to the vessel centerline; this is used to draw two-dimensional contour of the vessels (4). Data objects (centerlines, vessel lofts and trees, etc.) are shown in the data manager (5).

BCs are subsequently assigned, including upstream or downstream vascular reduced order models. The vessel walls can be modelled as rigid or made deformable via a coupled momentum method (Figueroa, et al., 2006), and spatially-varying mechanical properties of the arterial wall can be specified. External tissue support can also be specified, modelling the influence of nearby anatomical structures on the blood vessels. BC can be Neumann, Dirichlet, or based upon the coupled multidomain method (Vignon-Clementel, et al., 2006), which imposes a Dirichlet-to-Neumann relationship at the face. Boundaries can optionally use “backflow stabilization” (Hughes & Wells, 2005) to prevent numerical instability during periods of flow reversal at Dirichlet-to-Neumann surfaces; this critical feature is required to avoid the use of non-physiological “flow extensions” of the 3D domain often in other software packages. Numerical simulation parameters such as time stepping, and non-linear iteration strategies are then set within the GUI. Simulation files are written and passed to the parallel incompressible Flowsolver, which uses linear tetrahedral finite elements and a Streamline Upwind Petrov-Galerkin (SUPG) stabilized scheme for the spatial discretisation of the Navier-Stokes equations (Whiting & Jansen, 2001), and a second-order accurate generalized-α method (Jansen, et al., 2000) for time discretization. The Flowsolver scales efficiently to thousands of cores of HPC hardware, meaning that it can compute pressure and flow fields in very complex problems. Results are retrieved and can be visualized in the CRIMSON GUI. The GUI allows for field-based mesh refinement (Sahni, et al., 2006) and repetition of the analysis if necessary. Visualisation is also possible in third-party applications, such as Paraview (Paraview, 2016).

The CRIMSON Flowsolver consists of a core Fortran FEM solver, based on PHASTA (Sahni, et al., 2009) and closely related to that of SimVascular. We have made significant additions, including: (1) the conversion of the codebase from Fortran 77 to Fortran 90; (2) the addition of a modern C++11 layer to manage extensions, including the powerful new arbitrary BC design and control system; (3) the associated Python interface for dynamically adjusting BC parameters during simulations of transitional physiology; and (4) the use of the Verdandi libraries for data assimilation (Chapelle, et al., 2013). Additional modifications include the integration of the Google Test framework (Alphabet Inc., 2016), and our ongoing work towards simulation of non-linear vessel wall-blood flow interactions (Le Tallec, et al., 2001), (Nama, et al., Submitted) and flexible pipelines for simulation of systems of reaction-advection-diffusion (RAD) transport (Lynch, et al., 2020).

To assist with BC design, CRIMSON provides the Netlist Editor Boundary Condition Toolbox (NEBCT), shown in Figure 3. This presents a drag & drop interface for assembling LPNs, represented using electronic circuit symbols and invoking the standard analogy between fluidics and electronics. NEBCT allows tagging of components for Python-specified dynamic control during simulation and generates user-customizable Python controller class definitions, under the CRIMSON Control Systems Framework. NEBCT is based upon the QSapecNG electrical circuit simulation software (Arthurs & Figueroa, 2016) (Manetti, et al., 2012).

**Figure 3.**
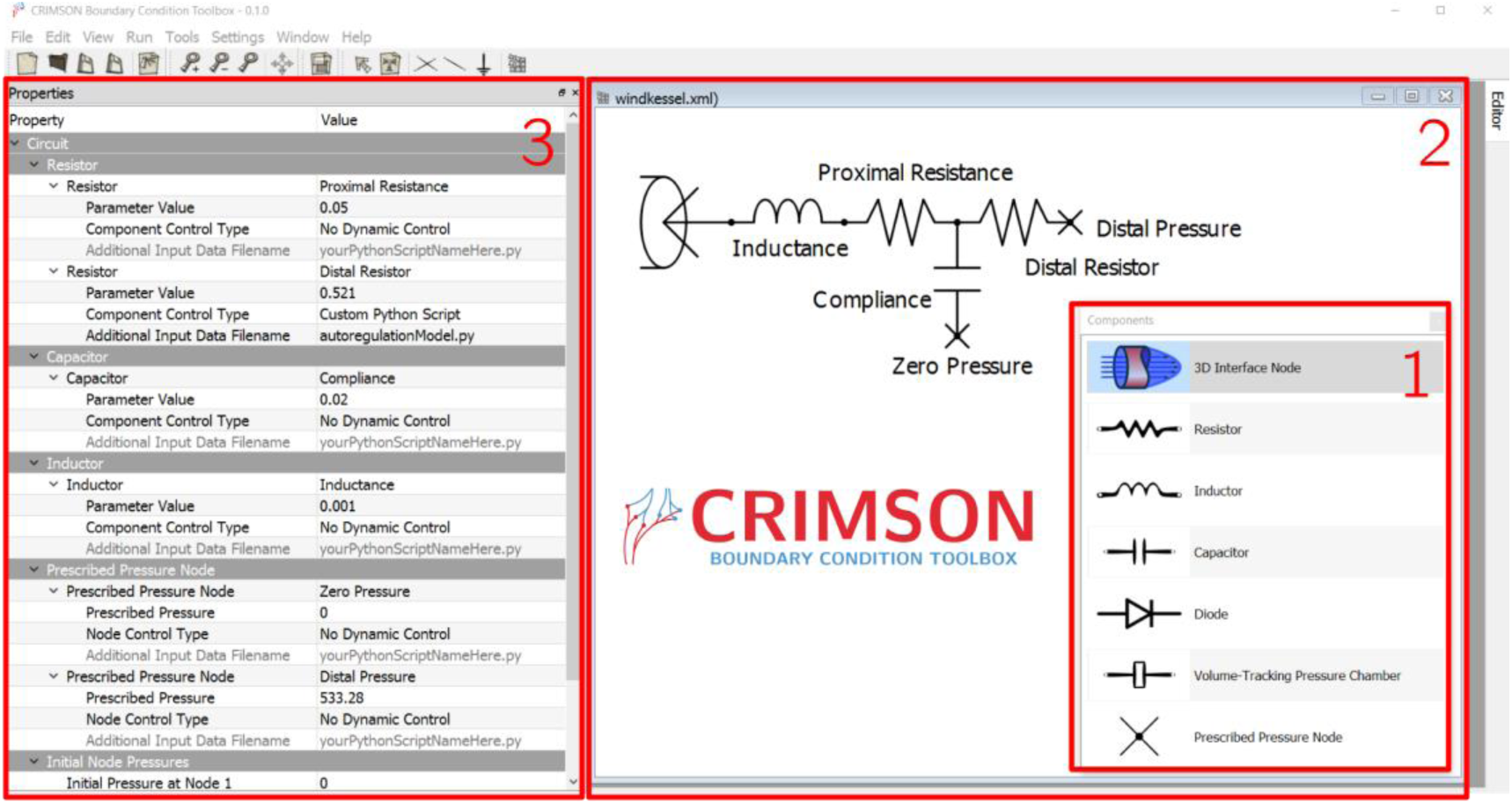
The CRIMSON Boundary Condition Toolbox. Researchers can use this tool to design custom LPN boundary condition models by choosing and arranging components from the toolbox (1) on the workplane (2). Parameters and other component properties can be set in the Properties pane (3). Here, a depiction of a Windkessel model is being created.

The CRIMSON GUI embeds another Python interpreter, which enables functionalities ranging from script-based specification of the local vessel wall thickness and elastic modulus, to definition of Python “Solver Setup” modules; this permits straightforward customization of the CRIMSON GUI’s interface widgets and output files, providing a pathway to compatibility with different fluid solvers. Thus, research groups may write their own Solver Setup to generate simulation files for their own simulation software; the procedure is documented online (https://crimsonpythonmodules.readthedocs.io/en/latest/). For example, a Solver Setup has been written for the CHeart fluid solver (Lee, et al. 2016) (Spazzapan, et al. 2018). Some additional details regarding the CRIMSON design and workflow have previously been presented (Khlebnikov, et al., 2016).

## Results

We demonstrate CRIMSON using three different examples, with all necessary input files provided in the supplement. We present two application examples (Cases A and B) highlighting specific aspects of CRIMSON’s capabilities, followed by an example demonstrating the entire workflow (Case C). In Case A, we demonstrate some of the fundamental features of CRIMSON, including running Navier-Stokes simulation in a patient-specific aortic model. In Case B, we demonstrate segmentation of PC-MRI data to create a patient-specific aortic inflow boundary condition. In Case C, we demonstrate the simulation of dynamic adaptations of blood flow in a surgical planning application.

### Example Case A: STL Geometry Import, Mesh Generation, Netlist Boundary Conditions, 3D Simulation on a Desktop PC

In Example Case A, we demonstrate the simulation workflow in the case where a geometric vascular model has been created using third-party software (Figure 4), which may be idealised or patient-specific. CRIMSON supports this via a solid model import feature. Note that additional boundary condition data (flow rates at the various model outlets) was assumed in this case; typically, additional patient data such as blood pressure or regional flow rates would be used to parameterize the boundary conditions.

**Figure 4.**
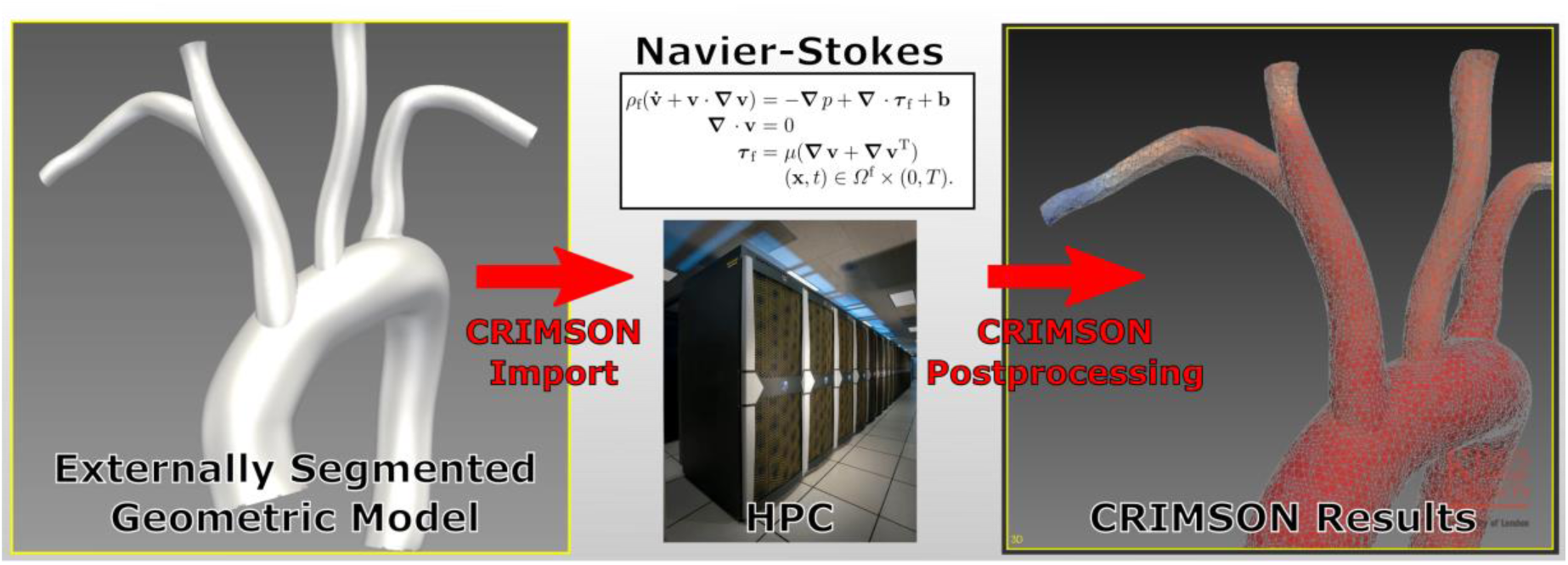
A simplified overview of CRIMSON’s ability to provide Navier-Stokes blood flow simulations in geometric models segmented from medical image data using third-party software. *HPC image: public domain. https://commons.wikimedia.org/wiki/File:Pleiades_row.jpg*

In addition to demonstrating how to work with imported geometries, this example highlights how to use the NEBCT to design simple LPN boundary conditions, how to run low-resolution 3D simulations on a desktop PC (or if the user wishes, high-resolution versions on HPC), how to refine the mesh after a simulation to focus computational effort in regions of high error in subsequent simulations, and how to visualise results within CRIMSON.

### Example Case B: Using Phase-Contrast MRI Data as a Patient-Specific Boundary Condition

A unique feature of CRIMSON is the ability to impose image-derived velocity profiles as boundary conditions. Typically, this is used at the aortic root, enabling study of the impact of valvular pathologies – which critically shape the inflow profile – on aortic haemodynamics. Figure 5 illustrates the concept. Imposing patient-specific inflow velocity profiles enables direct comparison with idealized (e.g. healthy) or hypothetical post-operative scenarios (e.g. following valve replacement procedures), without changing any of the other parameters of the system. It has been shown that CH which use idealised inflow profiles in cases where the patient’s valve is diseased can lead to significantly altered simulation results (Youssefi, et al., 2018) (Gomez, et al. 2018) (Youssefi, et al., 2017) (Youssefi, et al., 2016).

**Figure 5.**
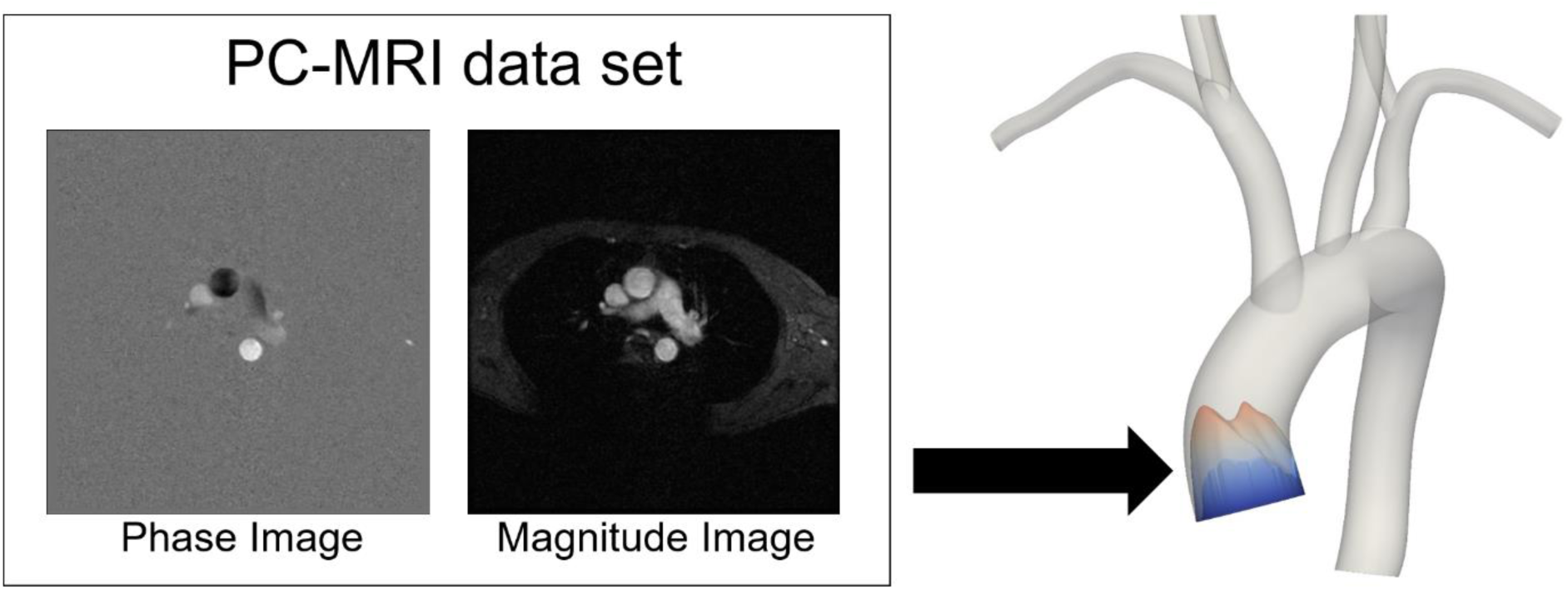
A PC-MRI (phase & magnitude images) plane slicing through the aortic root can segmented and processed using CRIMSON, so that patient-specific velocity profiles can be imposed as boundary conditions on the model.

The supplementary files in Example Case B show how to use CRIMSON to segment the provided PC-MRI dataset, and impose it upon the provided model.

### Example Case C: Cardiovascular Risk Assessment Post-Liver-Transplant

In Alagille syndrome patients in need of liver transplant, it is important to understand whether their cardiovascular system can withstand the stresses of the transplant surgery (Alagille et al. 1987, Emerick et al. 1999, Turnpenny & Ellard 2012, Kamath et al. 2010). A major concern is that the heart may not be able to support the demands placed on it during *post-reperfusion syndrome* (PRS) – the result of a sudden release of vasoactive mediators into the circulation after reperfusion of the new liver. PRS is characterised by a decrease in mean arterial pressure and systemic vascular resistance, and an increase in pulmonary pressure and central venous pressure (Aggarwal et al. 1987). These factors have the effect of simultaneously increasing the workload on the heart, and - due to the drop in mean arterial pressure - decreasing the coronary perfusion pressure gradient. Thus, the supply of oxygen to the ventricles may become insufficient, placing the patient at risk of cardiovascular complications.

To study this, a subject-specific model of an Alagille syndrome patient was created in CRIMSON, consisting of two (pulmonary and systemic) 3D domains segmented from magnetic resonance images, and a closed-loop circulatory system created using NEBCT (Figure 6). In the present article, we illustrate the difference between the myocardial oxygen supply and demand in one region of the left ventricular myocardium; for further details see Silva Vieira et al. 2018. This difference was computed using a model of myocardial oxygen supply and demand (Arthurs et al. 2016) and implemented within a coronary boundary condition using the CRIMSON Control Systems Framework.

**Figure 6.**
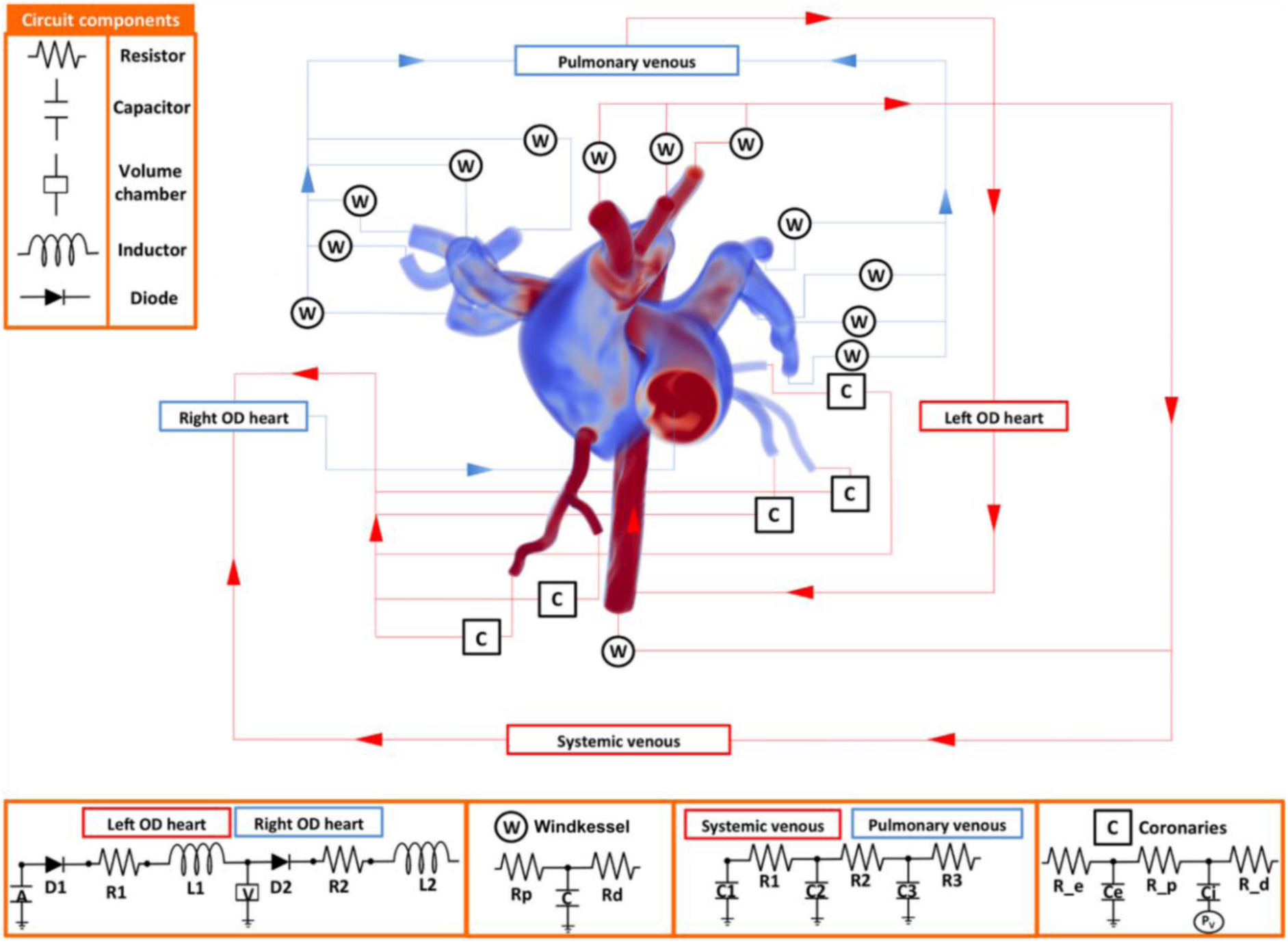
A complex, two-domain patient-specific 3D model with full closed-loop boundary condition system created in CRIMSON. This was used in a study of cardiovascular complications during liver transplant (Silva Vieira et al. 2018). Image used under CC BY 4.0 license.

Figure 7 compares the imbalance between myocardial oxygen demand and supply (“myocardial hunger”) in a baseline case (case C1), calibrated to patient haemodynamic data at rest, to that same imbalance under a simulated PRS condition (case C2). No long-term mismatch is detected under baseline conditions (the long-wave oscillation is a computational artefact as the simulation reaches a fully periodic state). Conversely, the continual growth in myocardial hunger under PRS indicates that this patient may be at risk of cardiovascular complications post-transplant.

**Figure 7.**
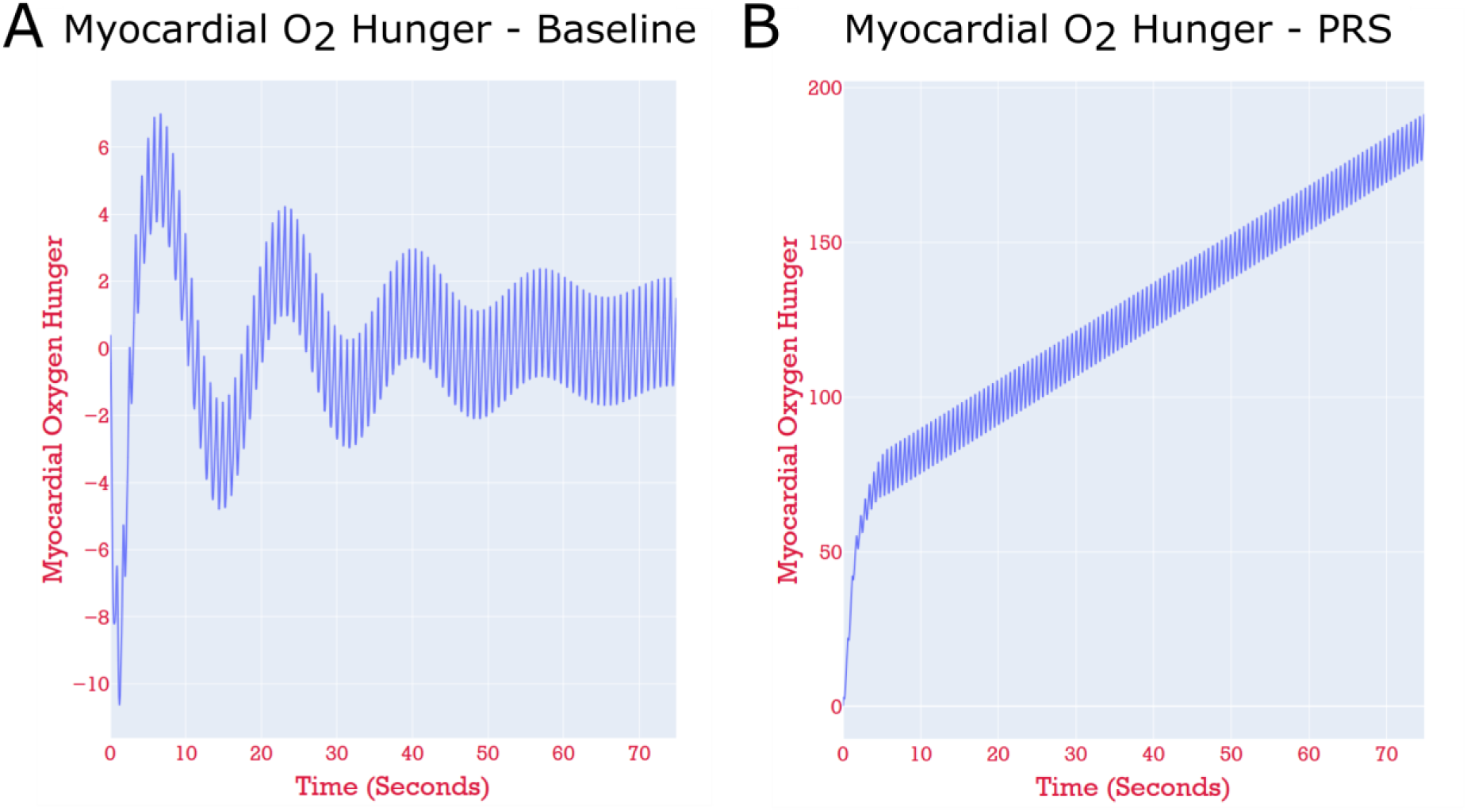
The results of running a pure zero-dimensional simulation of the model shown in Figure 6. Note the differing y-axis scales. **Panel A:** Case C1 - the time evolution of the myocardial hunger in the left anterior descending (LAD) perfusion territory of the myocardium, in the patient-specific simulation under baseline resting conditions. The blood supply is such that the hunger is kept close to zero for the entire simulation. **Panel B:** Case C2 - the model with parameters perturbed to simulate PRS. The hunger grows continually, indicating that the blood supply to this perfusion territory is insufficient, and thus, myocardial ischaemia.

The supplement describes how to run this case in 0D and visualize the results, as well as how to run in 3D, and visualize parameters such as local pressures and flows. The 0D simulation is presented here so that it is computationally tractable for most readers. Both the 0D and 3D simulations are presented in detail elsewhere (Silva Vieira et al. 2018). The closed-loop boundary condition circuit specified in this problem is visualized in Figure 8; guidance on how to perform this visualisation is also provided. Because running this model in 3D requires around 20,000 CPU core-hours, the input files provided in the supplement for Cases C1 and C2 are configured to run in a simplified 0D mode. Readers with access to HPCs can follow the instructions to enable full 3D simulation.

**Figure 8.**
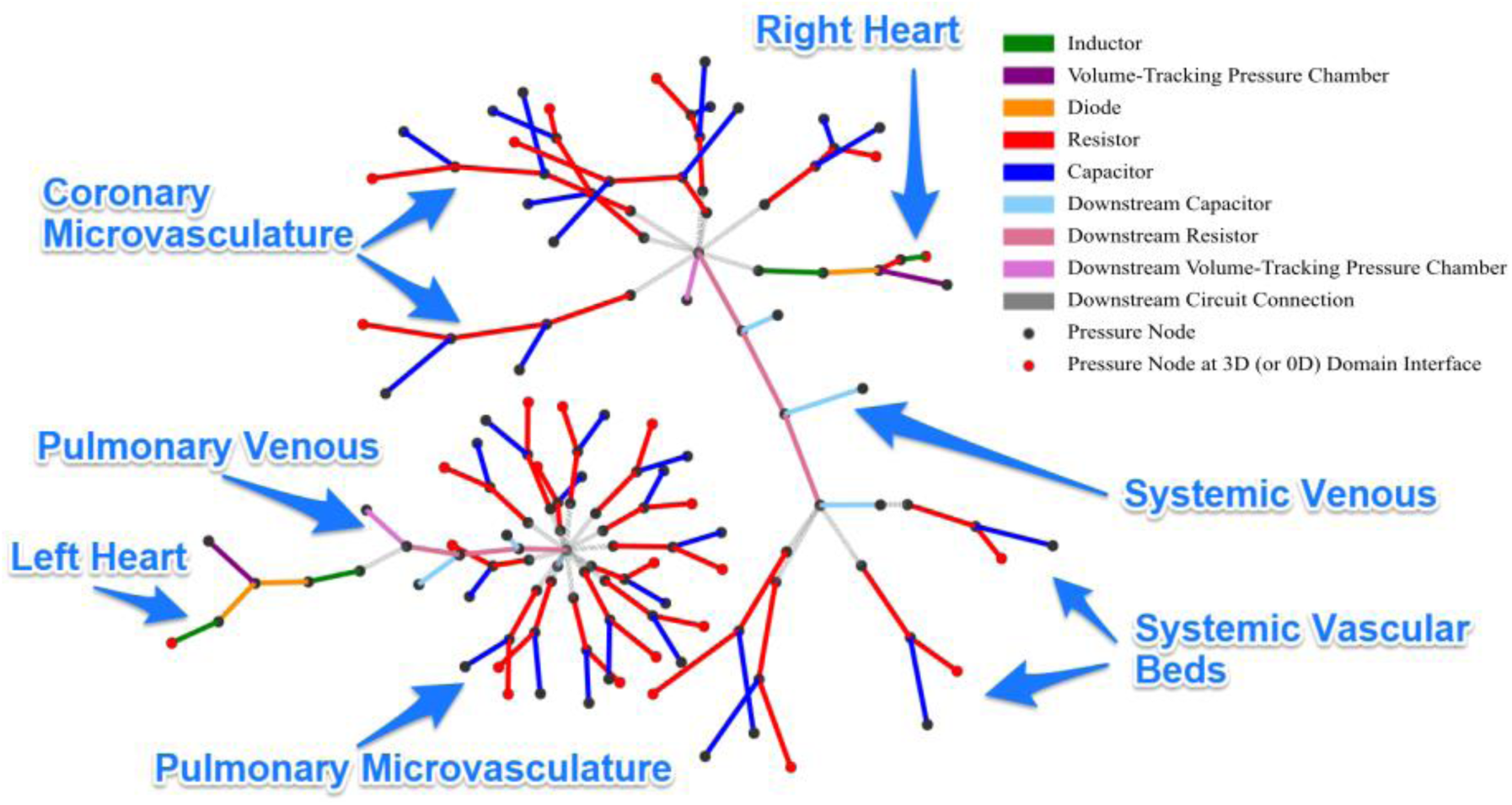
Visualisation of the Netlist boundary condition circuits shown in Figure 6. There are two connected components; during simulation, these two are joined by the two connected components of the 3D domain – the pulmonary arteries and the systemic arteries. Pressure nodes (between components) are shown as dots. Red dots indicate points of interface with the 3D domain. Lines indicate LPN components and are colored according to type (resistor, capacitor, etc.).

### Further applications of CRIMSON in the literature

Another use-case for CRIMSON is preoperative surgical planning. For example, CRIMSON was applied to a case of open surgical correction of a Fontan procedure, in a patient suffering from unilateral pulmonary arteriovenous malformations. After constructing a model reproducing the patient’s pre-operative haemodynamics, two post-operative models were created representing two possible surgical repairs. One was seen to be superior in terms of balancing hepatic venous flow between the pulmonary arteries, and the multidisciplinary surgical team chose this option for the patient (van Bakel, et al., 2018). Post-operative clinical data agreed with the simulated predictions.

In another study, CRIMSON was used to evaluate the suitability of different designs of a medical device – vascular endografts for aortic arch repair – in terms of their impact on patient haemodynamics. Evaluating design alternatives *in vivo* is neither practical nor ethical, so computational modelling is the only option. The results indicated that the different devices offered significantly different hemodynamic metrics in terms of blood shearing and blood flow to the brain (van Bakel, et al., 2018b).

CRIMSON has been used to systematically examine anatomical risk factors in type-B aortic dissection (TBAD) using idealized models. CRIMSON models permitted isolated changes to specific parameters present in TBAD, including vessel curvature, tear size and shape, false lumen location, and treatment state. The results indicated which combinations lead to unfavourable haemodynamic states, and agreed with clinical studies (Ben Ahmed, et al., 2016). A similar study employing a patient-specific model demonstrated that dissection imposes a significant additional workload on the heart, and that the number of additional communicating tears between the true and false lumen has a substantial impact on volumetric flow and peak pressure in the true lumen (Dillon-Murphy, et al., 2016).

In a study reproducing haemodynamics in a patient with a Blalock-Taussig (BT) shunt, CRIMSON was used to determine that the BT shunt’s diameter had reduced by 22% since implantation due to neointimal growth. Because the shunt size was below the MRI resolution, CRIMSON was critical in determining this diameter change (Arthurs, et al., 2017).

A patient-specific model involving three virtual interventions – open repair, conformable endografting, and fenestrated endografting – to repair a kinked ascending aortic graft in a patient-specific model was studied using CRIMSON (Nauta, et al., 2017). The results included evaluation of a metric of platelet aggregation potential, PLAP (Di Achille et al. 2014, Shadden & Hendabadi 2013), and indicated that surgical repair or the fenestrated endograft would most profoundly reduce thrombus formation risk.

One of the key novel aspects of CRIMSON is its tools for studying cardiovascular control systems and auto-regulations. These include the role of the baroreflex system in adjusting blood pressure after a change in body posture (Lau, et al., 2015), and how the coronary flow autoregulation system responds to changes in exercise intensity in a patient-specific model (Arthurs, Lau et al., 2016).

## Availability and Future Directions

We invite readers to download installers for Microsoft Windows from www.crimson.software for both CRIMSON GUI and CRIMSON Flowsolver. Alternatively, users can follow the documentation on the CRIMSON website, describing how to interface the GUI with their own fluid solver. The source code for the GUI is available on GitHub under GNU Affero General Public License v3.0 (https://github.com/carthurs/CRIMSONGUI), and for the Flowsolver under the GNU General Public License v3.0 (https://github.com/carthurs/CRIMSONFlowsolver). Both licenses further include the Commons Clause.

Figure 9 depicts several additional CRIMSON capabilities already available or currently under development. Panel A shows results obtained with a pre-existing custom aortic valve stenosis model, carried out using CRIMSON NEBCT and Control Systems Framework capabilities (Mynard, et al., 2012), which enabled straightforward implementation without requiring expert knowledge of the Flowsolver. Panel B shows results obtained using Lagrangian particle tracking analysis of haemodynamics in a “Zone 0” thoracic endovascular aortic repair (van Bakel, et al., 2018b). The color scale indicates PLAP (platelet activation potential), a metric used to quantify the shear rate experienced by massless particles advected by the flow. This index has been linked to risk of thrombus formation (Nauta, et al., 2017). Panel C shows the estimation of BC model parameters during the application of a sequential Reduced-Order Unscented Kalman Filter (ROUKF). The filter is used to determine suitable parameter values in agreement with time-resolved provided patient data, such as localised pressure or flow measurements (Moireau, et al., 2013, Chapelle, et al., 2013, Arthurs, et al., 2020). Panel D displays results of a volumetric mesh motion algorithm, which has been developed as a component of an Arbitrary Lagrangian Eulerian (ALE) based FSI framework utilizing a nonlinear shell formulation (Nama, et al., Submitted). This method will enable simulation of large vascular strains and motions, incorporating biologically-relevant constitutive models for the vessel wall, and involves a novel mathematical formulation that retains displacement-only degrees of freedom for the shell description for vessel wall, thereby ensuring minimal computational costs compared to other ALE methods. Panel E illustrates the application of a custom ODE model, composed in a few tens of lines of Python by the user, which at run-time can adjust the parameters of an attached BC component.

**Figure 9.**
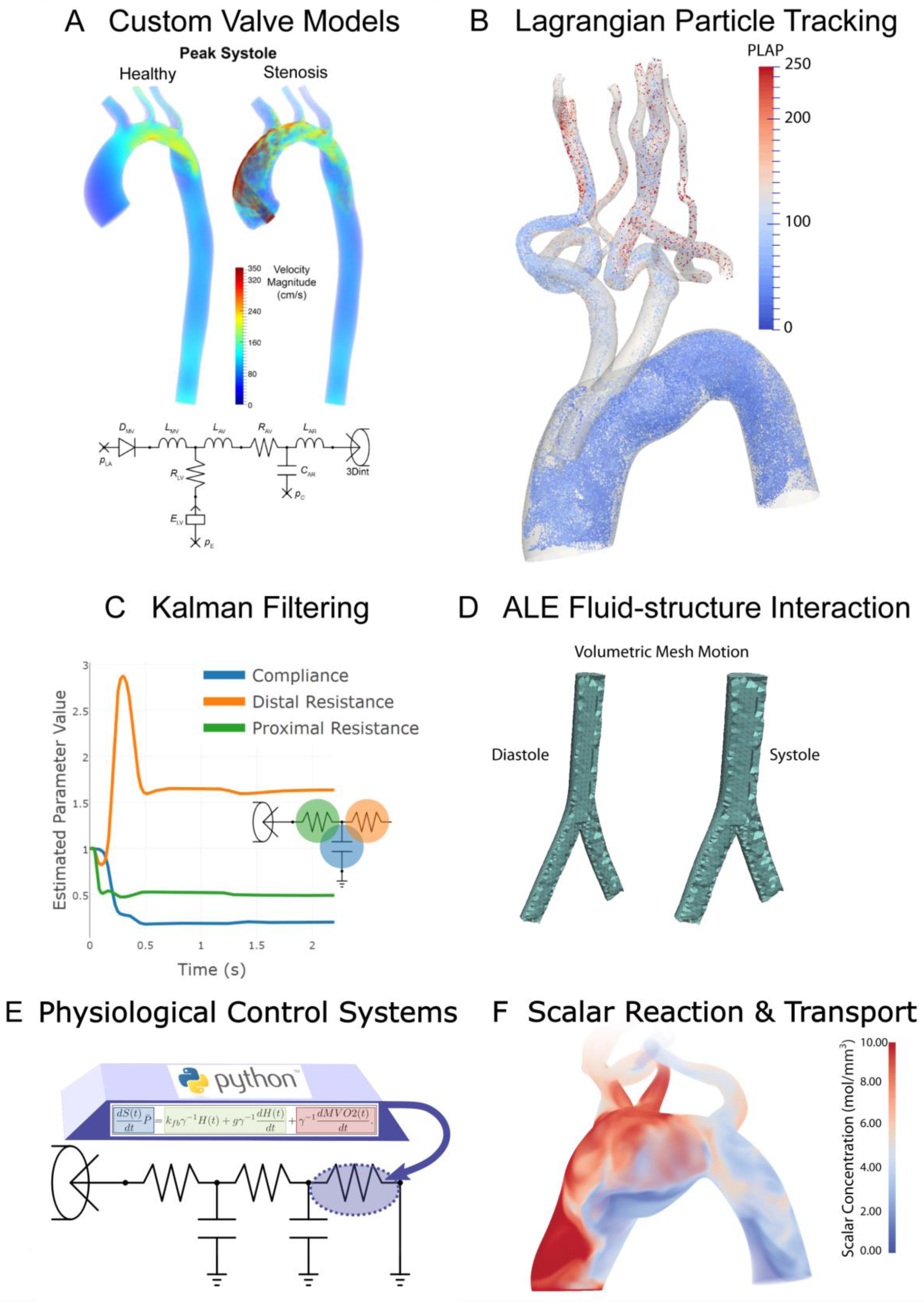
Examples of advanced and upcoming CRIMSON features. **A:** Custom aortic valve model implementation using the NEBCT and Python Control Systems Framework. **B:** Example of a Lagrangian particle tracking study of massless particle transport in the blood stream. **C:** Time history of parameter convergence during Kalman filter data assimilation. **D:** Arbitrary Lagrangian-Eulerian deformation of a vascular mesh between diastole and systole. **E:** Support for arbitrary run-time Python code for modelling cardiovascular control systems and changes of state, by controlling component parameters during simulations. **F:** Scalar reaction-advection-diffusion (RAD) problem with one species transported in the blood stream. Run-time specification of arbitrary reactions between tens of species is under current development.

Panel F illustrates the solution of RAD transport problems coupled with flow. Here, a single scalar with zero initial condition is considered, an advective flux is imposed at the inlet with a zero total flux condition on all walls and outlets. This allows simulation of transport under high Peclet numbers and interaction of scalar species, including proteins, drugs, contrast agents and heat, and will enable study of phenomena such as thrombogenesis (Lynch, et al., 2020). CRIMSON will support arbitrary numbers of reacting species, with reaction terms specified by the end-user at run-time in Python.

A further planned enhancement is the Integration of a one-dimensional (1D) incompressible Navier-Stokes solver, which enables simulating extensive vascular networks over many cardiac cycles in almost real-time, while retaining features such as pulse wave propagation phenomena (Xiao, et al., 2014). This 1D flow solver could be used in combination with graph theoretic approaches to accurately and efficiently model blood flow in disease conditions (Banerjee, et al., 2019).

We believe that CRIMSON is an enticing prospect for CH researchers and look forward to working with the community as we continue its development. We aim to be responsive to community input, via our support mailing list, and look forward to receiving feedback from our ever-expanding user-base on how CRIMSON can best support their work.

## Acknowledgements

We gratefully acknowledge support from the European Research Council under the European Union’s Seventh Framework Programme (FP/2007-2013) / ERC Grant Agreement n. 307532, the United States National Institutes of Health Grants U01 HL135842 and R01 HL105297, and the United Kingdom Department of Health via the National Institute for Health Research (NIHR) comprehensive Biomedical Research Centre award to Guy’s & St Thomas’ NHS Foundation Trust in partnership with King’s College London and King’s College Hospital NHS Foundation Trust, The Centre for Medical Engineering (CME) at King’s College London, and the Wellcome Trust Institutional Strategic Support Fund grant to King’s College London under grant number [204823/Z/16/Z]. JM is funded by the National Health and Medical Research Council of Australia and the National Heart Foundation of Australia.

## Supplementary Material

The version of CRIMSON GUI and CRIMSON Flowsolver current at the time of publication of this article is provided in the supplementary data package. However, a newer version may be available from www.crimson.software, as suggested in each of the cases below. To install CRIMSON, first run the CRIMSON-2019.11.01 exe installer, then run the CRIMSON_Flowsolver exe installer. The order is important. If you use the default installation paths in both, then the GUI will be able to find and run the flowsolver. If you install the GUI to a non-default location, you must change the Flowsolver’s install path in the same manner.

If you wish to build from source, follow the instructions in the readme.md file in the CRIMSON Flowsolver repository https://github.com/carthurs/CRIMSONFlowsolver; and analogously for the GUI repository https://github.com/carthurs/CRIMSONGUI.

### Running Example Case A

The tutorial pdf for this example case will guide you through the complete process of creating a model, running the simulation and examining the results.

#### File Manifest

1. **ExampleCaseA\ExampleA-Meshing-BC-Simulation-visualisation-refinement.pdf** – the instructions for running this case
2. **ExampleCaseA\DataFiles\Solid_model.stl** – the surface triangular mesh of the vascular geometry we will use
3. **ExampleCaseA\DataFiles\steady.flow** – specification file for a constant steady inflow at the aortic valve; for use with the Prescribed Velocities inflow boundary condition

#### Steps to Follow

1. Download and install CRIMSON from www.crimson.software, or from the installers in this data package. If using the installers in this data package, install *CRIMSON-2019*.*11*.*01-win64-trial*.*exe* first (the GUI), followed by *CRIMSON_Flowsolver_1*.*4*.*2_2019*.*11*.*01-installpath*.*exe* (the Flowsolver). If you adjust the installation path for the GUI, ensure you make the corresponding change for the flowsolver, so that the GUI can find the Flowsolver.
2. Open the document ExampleA-Meshing-BC-Simulation-visualisation-refinement.pdf from the ExampleCaseA folder of the Supplementary Files to this article. Follow the instructions therein.

### Running Example Case B

The supplementary pdf for this example will guide you through segmenting and imposing a patient-specific aortic inflow velocity profile from a provide PC-MRI dataset.

#### File Manifest

1. **ExampleCaseB\DataFiles\Patient-PCMRI.mitk** – the CRIMSON scene save file to start from, containing a geometric model to impose the PC-MRI boundary condition upon
2. **ExampleCaseB\DataFiles\PC-MRI_Images\Phase\** – a folder containing the PC-MRI phase images throughout one cardiac cycle
3. **ExampleCaseB\DataFiles\PC-MRI_Images\Magnitude\** – a folder containing the PC-MRI magnitude images throughout one cardiac cycle
4. **ExampleCaseB \ExampleB-PC-MRI-specificVelocityProfiles.pdf** – the instructions you should follow in order to impose a PC-MRI-derived inflow boundary condition on a model of the aorta

#### Steps to Follow

1. Download and install CRIMSON from www.crimson.software, or from the installers in this data package. If using the installers in this data package, install *CRIMSON-2019*.*11*.*01-win64-trial*.*exe* first (the GUI), followed by *CRIMSON_Flowsolver_1*.*4*.*2_2019*.*11*.*01-installpath*.*exe* (the Flowsolver). If you adjust the installation path for the GUI, ensure you make the corresponding change for the flowsolver, so that the GUI can find the Flowsolver.
2. Open the document ExampleB-PC-MRI-specificVelocityProfiles.pdf from the ExampleCaseB folder of the Supplementary Files to this article. Follow the instructions therein.

### Running Example Case C

The supplementary pdf for this case will guide you through running a simulation of a patient under rest conditions, and then of the same patient under post-liver-transplant conditions. Many parameters of the haemodynamics could be examined and compared between the two, but in this case we choose to examine the oxygen supply/demand balance in the myocardium under these two states, and will observe that coronary perfusion is insufficient in the post-operative simulation.

#### File Manifest

1. **ExampleCaseC\Mitk_Scenes** for the Resting and PRS conditions – these are the CRIMSON saved state files
2. **ExampleCaseC\AdditionalInputFiles** – Files prepared for this specific case (such as custom CRIMSON Control Systems Toolbox Python scripts, and loop-closing circuits, which have to be made manually outside the GUI). These are for combining with the CRIMSON GUI output for each case to create complete simulation input files
3. **ExampleCaseC\ReadyToUseFiles** – folders containing simulation input files in a state which can be run immediately. These files are configured to run in purely zero dimensions, and essentially consist of a combination of the CRIMSON Solver Setup output from the MITK scenes, and the AdditionalInputFiles. You will only need these if you do not wish to generate your own simulation files, as is one of the two options described in ExampleC-simulating-Rest-vs-PRS-in-Alagille-patients.pdf
4. **ExampleCaseC\ExampleC-simulating-Rest-vs-PRS-in-Alagille-patients.pdf** – the instructions you should follow in order to create and run the two simulation states for the patient.

#### Steps to Follow

1. Download and install CRIMSON from www.crimson.software
2. Open the document ExampleC-simulating-Rest-vs-PRS-in-Alagille-patients.pdf from the ExampleCaseC folder of the Supplementary Files to this article. Follow the instructions therein.

#### Libraries and Frameworks used in CRIMSON

The following pre-existing libraries are integral to the CRIMSON software.

The Medical Imaging Interaction Toolkit (MITK) (Wolf, et al., 2005) was chosen as the foundation for the CRIMSON GUI due to its established position as an open-source package for development of interactive medical image processing and ongoing development by the German Cancer Research Center. It provides functionality for working with medical image datasets, including multiple orthogonal and 3D views of the image volume, data storage and undo/redo functionality, and image processing and segmentation algorithms (Nolden, et al., 2013) via the Insight Toolkit (ITK), Visualization Toolkit (VTK) and Qt (The Qt Company, 2016). Additionally, it includes a Python interface for easy prototyping and MITK release 2016.11 integrates VMTK (The Vascular Modeling ToolKit, Antiga et al. 2008), a reference package for vascular segmentation and automatic centreline extraction. MITK is well-regarded, providing a foundation for other research packages including by the more general GIMIAS (Larrabide, et al., 2009).

MITK itself is built upon the widely used visualisation tools of VTK, adopts segmentation tools from ITK, and is highly customizable due to the modular, cross-platform MITK BlueBerry framework and CTK (*the Common Toolkit*). MITK uses the Qt application framework to manage events and connect user interface (UI) elements, the MITK DataStorage system to manage state data (The Qt Company, 2016). Qt, together with the Python Qt wrappers, allows advanced users to add, remove or modify the widgets presented by the GUI when writing custom Solver Setups (see later in this section). The functionality for lofting or sweeping to create the solid model from the segmented vessel contours, blending the intersecting vessels into a full geometry, and important filleting and Boolean operations is provided by OpenCascade (OpenCascade, 2016). Volume meshing is provided by the TetGen (Si, 2015) library; optionally, this can be replaced with the high-quality commercial Simmetrix MeshSim library (Simmetrix, 2016).

*OpenCascade* (Open Cascade SAS, 2017) is an open-source C++ class library designed for rapid production of complex Computer Aided Design (CAD) applications. This class library makes it possible to define analytical models whose surfaces are parametrized by functions such as non-uniform rational basis splines (NURBS).

*Verdandi* is an open-source generic C++ library for data assimilation, developed by INRIA (Chapelle, et al., 2013). Data assimilation is a process that enables combining different sources of information to estimate the state of a dynamical system. By extension, this approach can be used to estimate parameters of a computational model if time-resolved data is available. Verdandi was designed so that computational models implemented in C++, Fortran, or Python could be make use of Verdandi via either a C++ or a Python interface.

*QSapecNG* (Manetti, et al., 2012) is an open-source program based on Qt for drawing linear analogue circuits and performing analysis. We modified it to create a tool for designing custom circuits for BC models (the CRIMSON Netlist Editor Boundary Condition Toolbox), and outputting them in a format readable by the CRIMSON Flowsolver.

## Notes

https://drive.google.com/file/d/1kl0PHcwRClzUuyLOPzxRNqE_h8_TpeFQ/view?usp=sharing

### Competing Interest Statement

CJA and CAF have a joint interest in CRIMSON Technologies LLC, a supporting company for CRIMSON, providing software and services to researchers. This is in addition to the open-source version of CRIMSON which is freely available.

